# Quencher-free fluorescence monitoring of G-quadruplex folding

**DOI:** 10.1101/2024.01.31.578026

**Authors:** Zach Parada, Tanner G. Hoog, Katarzyna P. Adamala, Aaron E Engelhart

## Abstract

Guanine-rich sequences exhibit a high degree of polymorphism and can form single-stranded, Watson-Crick duplex, and four-stranded G-quadruplex structures. These sequences have found a wide range of uses in synthetic biology applications, arising in part from their structural plasticity. High-throughput, low-cost tools for monitoring the folding and unfolding transitions of G-rich sequences would provide an enabling technology for accelerating prototyping of synthetic biological systems and for accelerating design-build-test cycles. Here, we show that unfolding transitions of a range of G-quadruplex-forming DNA sequences can be monitored in a FRET-like format using DNA sequences that possess only a single dye label, with no quencher. These quencher-free assays can be performed at low cost, with both cost and lead times ca. 1 order of magnitude lower than FRET-labeled strands. Thus, quencher-free secondary structure monitoring promises to be a valuable tool for testing and development of synthetic biology systems employing G-quadruplexes.

## Introduction

Guanine-rich sequences exhibit an unusual degree of polymorphism among nucleic acids.^1,2^ In addition to their single-stranded form and Watson-Crick duplex, G-rich sequences can exist in equilibria with four-stranded structures called G-quadruplexes, which form via Watson-Crick-Hoogsteen interactions between quartets of four guanines, which in turn coordinate a central cation. G-quadruplex (G4) structures have been employed in a range of synthetic biology applications, including sensing of diverse ligands for readout of metabolic states,^3–6^ forming nucleic acid-based nanomachines,^7,8,^ and biomimetic cofactor binding that enables catalysis of a wide range of electron transfer reactions.^9–12^ Monitoring the polymorphism and folding states of G-rich sequences is key to understanding their function. A variety of techniques have been employed for such monitoring, including high-resolution structural techniques such as NMR and crystallography,^13–16^ as well as higher-throughput spectroscopic techniques, including circular dichroism and FRET.17,18 Of these, FRET is among the most widely used, due to its experimental expediency and high throughput.^19^

In a typical execution, FRET experiments employ a fluorophore as a donor, and a second dye as an acceptor – typically one with low fluorescence quantum yield, which acts as a quencher. These experiments require the synthesis of dual-labeled probes. These probes have high associated cost, greater lead time, and low yields arising from the need to incorporate both 3□ and 5□ modifications and perform HPLC purification. For example, the current price of a 25 nt dual-labeled 5□-fluorescein/3□-BHQ-1 labeled oligonucleotide at 100 nmol scale is $265 ($26.50/guaranteed nmol) with an associated lead time of 2 weeks.

While performing FRET characterization of a switchable G4-based nucleic acid electron transfer catalyst,^11^ we serendipitously observed that a singly-labeled G-rich oligonucleotide provided FRET-like curves that acted as a reporter of strand folding state, with the G4 state of a singly labeled strand giving high fluorescence, and the single-stranded (ss) state giving low fluorescence. The use of singly modified dye-labeled oligonucleotides would greatly simplify and lower cost in experimental workflows. These are far more readily prepared than dual labeled oligonucleotides: a 25 nt 5□-fluorescein-labeled oligonucleotide at the same scale costs only $99.50 ($2.84/guaranteed nmol) with an associated lead time of 2 days. This ca. 1-order of magnitude reduction in both cost and lead time would greatly enable rapid prototyping and design-build-test cycles. Motivated by this, we sought to investigate whether the secondary-structure dependent fluorescence of singly labeled oligonucleotides we observed was a general phenomenon.

## Results and Discussion

We characterized a range of oligonucleotides capable of forming G4 structures by both fluorescence and UV absorbance **(Figure 1)**. As a benchmark, we initially examined the fluorescence and UV absorbance thermal transitions of **FAM-G4-IB**, a G4-forming oligonucleotide with a 5□-fluorescein and 3□-Iowa Black quencher **(Figure 1a-b, Supplementary Table 1)**. As expected, **FAM-G4-IB** gave a cooperative transition, indicative of a G-quadruplex (G4) to single-strand (ss) transition. In the absorbance trace, a decrease in absorbance at 295 nm was observed, consistent with the unfolding of a G4. In the fluorescence trace, the G4 state gave low fluorescence, and the ss state gave high fluorescence, consistent with the closer positioning of the 5L dye label and 3L quencher in space in the folded G4 state relative to the unfolded ss state. Both fluorescence and absorbance traces gave TM values that agreed with one another, consistent with both spectroscopic techniques reporting on the same unfolding transition in potassium buffer **(Figure 1b, Supplementary Tables 2 and 3)**, as well as sodium buffer **(Supplementary Figure 1a)**

**Figure 1.**
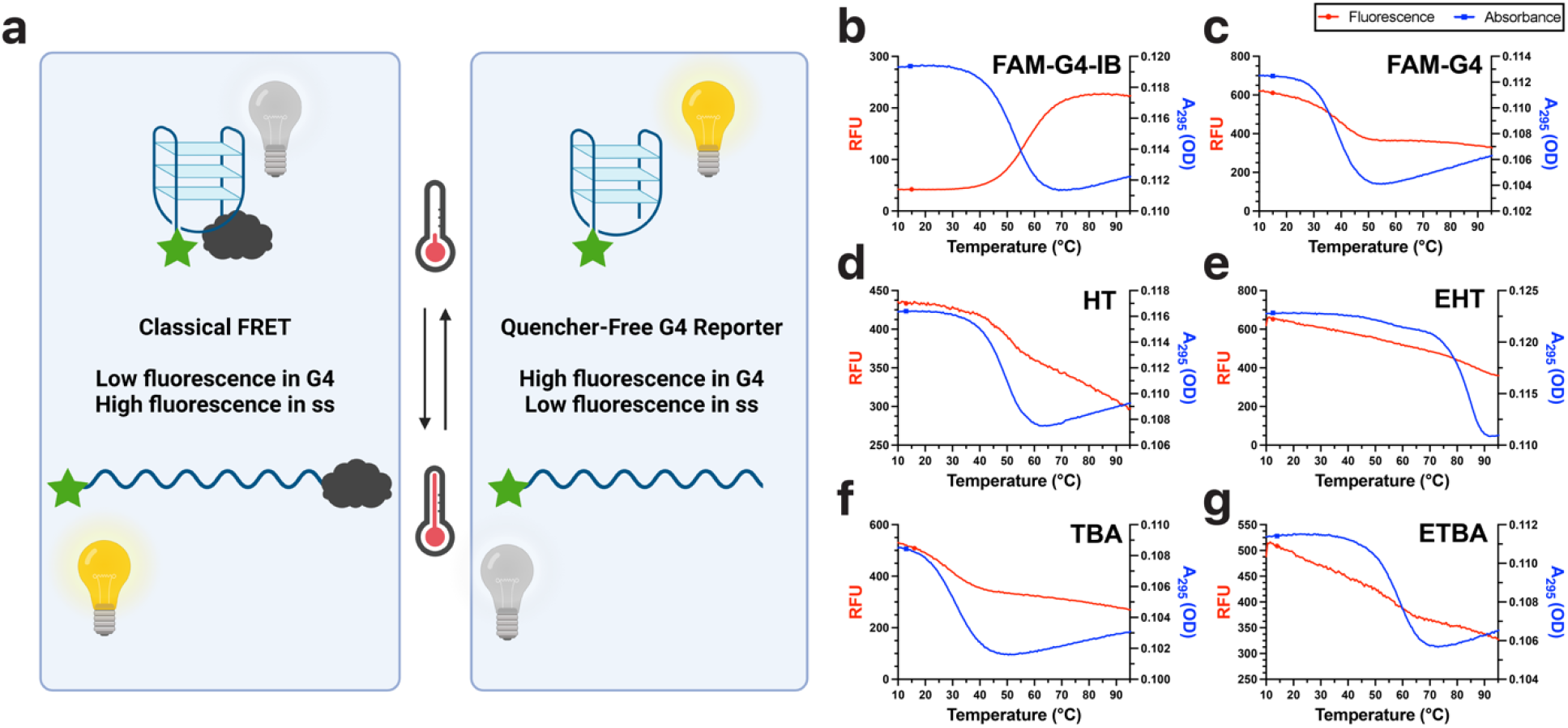
Quencher-free-fluorescence and absorbance-monitored thermal denaturation profiles both report on unfolding of various G-Quadruplex-forming sequences. (a) Graphical comparison of classical FRET and quencher-free fluorescence monitoring of unfolding transitions, (b) **FAM-G4-IB** with fluorescein and Iowa Black quencher, (c) **FAM-G4** with only fluorescein modification, (d) **HT** with four repeats of human telomere sequence, (e) **EHT**, a variant of **HT** with an expanded G-tract. (f) **TBA**, the 15 nt thrombin binding aptamer, (g) ETBA, a variant of TBA with an expanded G-tract. Cooperative transitions were observed in all cases, indicating G-quadruplex to single-stranded transitions. All data shown was collected in K^+^-containing buffer.

We next examined **FAM-G4**, the same sequence as **FAM-G4-IB**, with only a 5□-fluorescein end label and no 3□-quencher. This sequence also gave a quencher-free-fluorescence melting profile that corresponded with that observed by absorbance spectroscopy (K^+^ : **Figure 1c**, Na^+^: **Supplementary Figure** 1b). The fluorescence trace for **FAM-G4** trace was the inverse of that observed for **FAM-G4-IB**, with **FAM-G4** giving high fluorescence in the G4 state and low fluorescence in the ss state. The unfolding transition of FAM-G4-IB was stabilized relative to FAM-G4 (56.9 °C vs 38.6 °C in potassium solution, **Supplementary Table 2**) consistent with previous reports of stabilizing interactions associated with oligonucleotide labels. ^20^

We examined a suite of other G4-forming oligonucleotides. The G4 formed by four repeats of the human telomere sequence d(TTAGGG) (**HT**, K^+^ : **Figure 1d**, Na^+^: **Supplementary Figure 1c**) ^21^ exhibited the same behavior, with good agreement between absorbance and quencher-free-fluorescence traces.. An expanded variant of this sequence with repeats of d(TTAGGGG) (**EHT**, K^+^ : **Figure 1e**, Na ^+^ : **Supplementary Figure 1d**) also showed similar behavior.

We next sought to examine an aptamer sequence, as G-quadruplex structures are often found in this important class of functional RNA.22 The thrombin binding aptamer (**TBA**, K^+^ : **Figure 1f**, Na ^+^ : **Supplementary Figure 1e**), ^23^ which contains a G4 with only two guanine quartets, is known to fold stably in potassium, but not sodium solution. As expected, quencher-free-fluorescence and absorbance spectra reflected the presence of a cooperative unfolding transition in potassium solution only. An expanded version of this aptamer with a three-guanine quartet core (**ETBA**, K^+^ : **Figure 1g**, Na ^+^ : **Supplementary Figure 1f**) gave the expected increased stabilization relative to **TBA** and concurrence between quencher-free-fluorescence and absorbance traces as well. We observed agreement between quencher-free-fluorescence and absorbance traces for all G4-forming sequences tested (**Supplementary Tables 2** and **3**).

Guanine possesses the lowest redox potential of the canonical nucleobases, ^24,25^ and due to this, it is well-known to nonspecifically quench fluorescence by a mechanism that is thought to involve photoinduced electron transfer. Because of this, it is typically not recommended to have a guanine residue immediately adjacent to a dye label. The sequence we initially tested possessed no spacer between the dye and G4-forming sequence (K^+^: **HT, Figure 2a/TBA, Figure 2b**, Na^+^: **HT, Supplementary Figure 2a/TBA, Supplementary Figure 2b**). As expected, this gave the lowest fluorescence of the sequences tested, with fluorescence suppressed by 30-50% relative to labeled oligonucleotides with no spacer residue. Despite this, it still could report on unfolding, consistent with the formation of a G4 structure relieving guanosine-induced fluorescence quenching.

**Figure 2.**
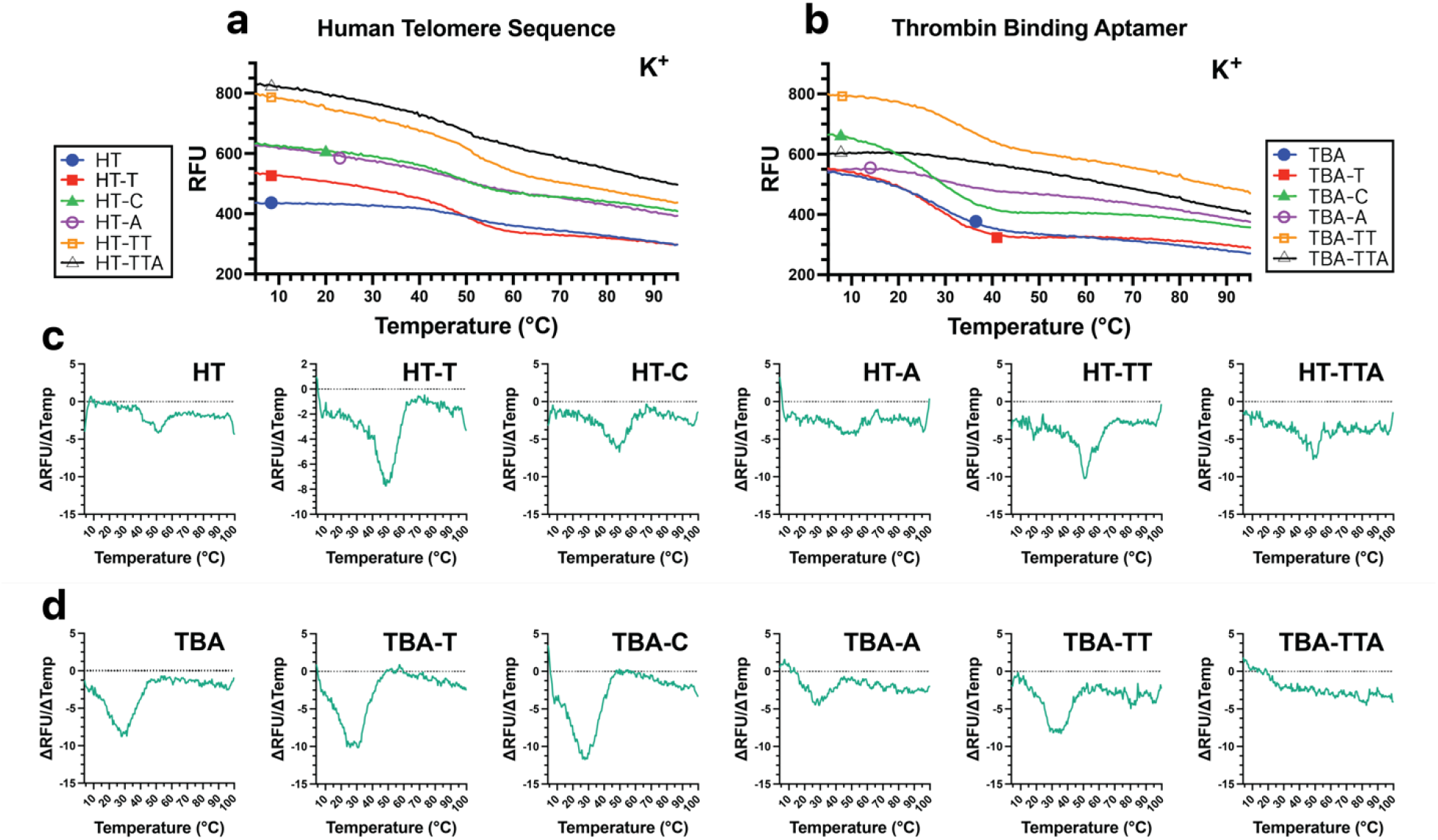
Influence of fluorophore proximity and spacer sequences on fluorescence transitions in two G-Quadruplexes. (a) Fluorescence thermal melts of human telomere sequence (HT) with zero-, single-, or multiple-nucleotide spacers. (b) Fluorescence thermal melts of thrombin binding aptamer (TBA) with zero-, single- or multiple-nucleotide spacers. (c) First derivatives of HT fluorescence vs temperature with single or multiple nucleotide spacers (d) First derivatives of TBA fluorescence vs temperature values with single or multiple nucleotide spacers. All data shown was collected in K+-containing buffer.

We sought to investigate whether the secondary structure-dependent effect we observed was dependent on the presence of “spacer” nucleotide residues between the 5□-dye label and the sequence being interrogated for folding activity. Accordingly, we prepared analogues of the **HT** and **TBA** sequences with variable spacers between the 5□-fluorescent residue and the G4-forming sequence of interest (**Figure 2**). We examined sequences containing a spacer consisting of each non-guanosine nucleotide: thymine (**HT-T/TBA-T**), cytosine (**HT-C/TBA-C**), and adenine (**HT-A/TBA-A**). Each of these sequences gave higher fluorescence values than the spacer-free sequences **HT** and **TBA**, with similar capacity to report on unfolding transitions (K^+^: **Figure 2**, Na^+^: **Supplementary Figure 2**).

Finally, we examined sequences containing longer dinucleotide (d(TT), **HT-TT/TBA-TT**) or trinucleotide (d(TTA), **HT-TTA/TBA-TTA**) spacers. Here, **TBA-TT** and **HT-TT** gave better performance as reporters than **TBA-TTA** or **HT-TTA**. In the case of d(TTA)-spacer containing oligonucleotides, only the potassium form of **TBA-TTA** gave a clear transition by fluorescence (K^+^: Figure 2, Na^+^: Supplementary Figure 2).

To investigate the impact of folding state on quenching behavior, we examined **TBA** and **HT** by circular dichroism. Consistent with our other results reported here, as well as prior reports of the unlabeled version of this strand, only the potassium form of **TBA** was folded into a G-quadruplex, while **HT** was folded in both potassium and sodium solution (Supplementary Figure 3). **HT** gave cation-dependent topologies, with the sodium form exhibiting spectra consistent with an antiparallel topology, and the potassium form exhibiting a hybrid topology. Thus, the different three-dimensional orientation of the sodium and potassium forms of **HT** could explain in part the different degrees of quenching phenomena observed in **HT-TTA** in different buffers. This strand shows an approximation of the limit of spacer length for quencher-free fluorescence monitoring of G4 folding. Thus, these results illustrate that considerable sequence flexibility exists as evidenced by the strands containing A, T, and C spacers, but that the shortrange phenomenon we report here is exquisitely distance-dependent and thus best employed for reporting on local folding and unfolding transitions.

To further verify that the observed fluorescence transitions resulted specifically from G4 formation and denaturation, we compared another G-quadruplex-forming human telomere sequence **HT-2** with its isomer **HT-2-I**, which was scrambled to disrupt G-tracts and suppress G-quadruplex formation (**Supplementary Figure 4**). **HT-2** was selected because it contained an equal number of G and non-G residues, enabling generation of the scrambled sequence **HT-2-I** with no consecutive guanine residues. We used **HT-2** Quencher-free fluorescence and absorbance monitoring of thermal melts of **HT-2** showed unfolding of G-quadruplexes, as expected. Despite identical guanine content in the two sequences, **HT-2-I** did not exhibit transitions in quencher-free fluorescence or UV-vis absorbance thermal melts of **HT-2**, consistent with the requirement for G-tracts for quenching in the single-stranded state, and for folding into a G-quadruplex structure to relieve this quenching.

Guanosine-mediated fluorescence quenching has been employed to monitor a range of phenomena, including hybridization, DNA processing, ribozyme kinetics, and nucleotide cleavage. ^26–32^ The results presented here extend the utility of this quenching phenomenon to a monitoring of the folding state of a G-quadruplex-forming strand. As G-quadruplex secondary structures continue to find increasing utility in a wide range of synthetic biology applications, we anticipate that the low-cost, high-throughput detection method afforded by quencher-free fluorescence monitoring of these unfolding transitions will provide an enabling technology for accelerating design-build-test cycles in these and other applications.

Our results also extend the literature on guanine-mediated single electron transfer reactions. The propensity of guanosine for oxidation has been invoked to explain the excited-state quenching observed when guanine nucleobases are present immediately adjacent to a fluorophore, as well as the sequence specificity associated with DNA damage and conductive behavior within DNA. ^33^ Our results illustrate that guanine-associated quenching is influenced by nucleic acid secondary structure. Thus, they suggest that other single-electron transfer reactions in which guanine acts as an electron donor could be similarly influenced by secondary structure.

## Materials and Methods

### Buffers

All experimental data were obtained using a 50 mM Li-HEPES buffer at pH 7.4. Each buffer additionally contained an alkali chloride salt. The lithium-containing buffer contained 50 mM LiCl (Sigma-Aldrich 203637), sodium-containing buffer contained 50 mM NaCl (Sigma-Aldrich S7635), and potassium-containing buffer contained 20 mM KCl (Sigma-Aldrich P9333).

### DNA Sample Preparation

All DNA oligonucleotides were procured from Integrated DNA Technologies as desalted grade (unlabeled or singly labeled) or HPLC purified (dual-labeled). Oligonucleotides were resuspended in water to prepare 100 µM stocks. Individual oligonucleotide stocks were further quantified through UV absorption at 260 nm and diluted to 50 µM sub-stocks for subsequent use; these stocks were stored at -20°C. Upon dilution into the appropriate buffer and prior to use, all samples underwent pre-folding by heating to 95°C for 5 minutes, followed by cooling to room temperature over a 3-hour period. Fluorescence and UV-vis thermal melts contained 5 µM 5’
s-fluorescein labeled oligonucleotide, while circular dichroism samples contained 20 µM 5’fluorescein-labeled oligonucleotide. Sequences for oligonucleotides employed are given in **Supplementary Table 1**.

### Fluorescence-Monitored Thermal Melts

Fluorescence measurements were performed using a Cary Eclipse fluorescence spectrophotometer (Agilent Technologies) equipped with a Peltier thermostatted multicell holder (Agilent Technologies). The excitation wavelength was set to 495 nm, and the emission wavelength was set to 520 nm. Excitation slits were set to 2.5 nm and emission slits were set to 5 nm. The photomultiplier tube (PMT) voltage was set to 565 V. After pre-folding (see **DNA Sample Preparation**), all samples underwent a temperature ramp from 4 °C to 100 °C at a rate of 0.5°C/min to minimize hysteresis. This was followed by a one-minute hold at 100°C, a subsequent cooling phase from 100 °C to 4 °C at the same rate, and an additional one-minute hold period at 4°C. Fluorescence measurements were taken at 0.5°C intervals, with a signal averaging time of 0.25 seconds. Repeating this experimental sequence allowed for a total of two heat/cool cycles.

### UV-Vis Monitored Thermal Melts

Thermal denaturation experiments were performed using a Cary 60 UV-Vis spectrophotometer (Agilent Technologies), equipped with a qCHANGER 6 temperature-controlled linear cuvette changer (Quantum Northwest) controlled by a TC 1/Multi temperature controller (Quantum Northwest). Liquid cooling for the Peltier device was provided by a EX2-755 liquid cooling system (Koolance). The above instruments were interfaced using Cary ADL scripts (Quantum Northwest) that allowed monitoring of absorbance vs. temperature at 295 nm. This allowed monitoring of G-quadruplex folding transitions. After pre-folding (see DNA Sample Preparation), samples were heated from 10 °C to 100 °C and then cooled from 100 °C to 10 °C, using 0.5 °C steps for each ramp. At each step, a 30-second hold was performed before each reading with a signal averaging time of 0.25 seconds. This protocol gave approximately the same ramp times as that for the fluorescence experiments described above.

### Circular Dichroism Spectroscopy for G-Quadruplex Topologies

Circular Dichroism (CD) measurements were carried out using a J-815 spectropolarimeter (JASCO) equipped with a Peltier temperature controller (JASCO). CD spectra were recorded between 200 nm and 600 nm, with continuous scanning mode and a scanning speed of 100 nm/min. The data integration time was fixed at 4 seconds per data point, and a data pitch of 1 nm and 2 nm bandwidth.

### Data Analysis

Data from fluorescence and absorbance melting experiments were imported using MATLAB 2023b software (MathWorks). Subsequently, individual traces were plotted, and thermal midpoints were identified utilizing Igor Pro version 9 software (WaveMetrics). These midpoints were determined as the inflection points of sigmoidal fits corresponding to each respective heating or cooling transition. The recorded thermal midpoints are provided in **Supplementary Tables 2** and **3**.

Circular dichroism data was imported directly into Igor Pro version 9 (WaveMetrics) for processing and graphical representation.

