## Supplementary Information for "Quencher-free fluorescence monitoring of G-quadruplex folding"

for

**Contents**

**Supplementary Table 1.** Oligonucleotides used in this work

**Supplementary Table 2.** Fluorescence melting temperatures of oligonucleotides

**Supplementary Table 3.** Absorbance melting temperatures of oligonucleotides

**Supplementary Figure 1.** Quencher-free-fluorescence and absorbance-monitored thermal denaturation profiles both report on unfolding of various G-Quadruplex-forming sequences in sodium solution.

**Supplementary Figure 2.** Influence of fluorophore proximity and spacer sequences on fluorescence transitions in various oligonucleotide G-tracts in sodium solution.

**Supplementary Figure 3.** Circular dichroism spectra of G4-forming oligonucleotides.

**Supplementary Figure 4.** Experiments with two isomeric sequences illustrate that quencher-free fluorescence reports on secondary structure, rather than sequence content.

### Supplementary Table 1. Oligonucleotides used in this work.

| **Sequence Name** | **Description** | **Sequence (5ʹ-3ʹ)** |
| --- | --- | --- |
| **FAM-G4-IB** | FAM- G4 Sequence - Iowa Black | /56-FAM/TGG GTT AGG GAA TTC GGG TTA GGG/3IABkFQ/ |
| **FAM-G4** | FAM - G4 Sequence | /56-FAM/TGG GTT AGG GAA TTC GGG TTA GGG |
| **HT** | FAM-Human Telomere | /56-FAM/GGG TTA GGG TTA GGG TTA GGG |
| **HT-T** | FAM-Human Telomere - T spacer | /56-FAM/TGG GTT AGG GTT AGG GTT AGG G |
| **HT-C** | FAM-Human Telomere - C spacer | /56-FAM/CGG GTT AGG GTT AGG GTT AGG G |
| **HT-A** | FAM-Human Telomere - A spacer | /56-FAM/AGG GTT AGG GTT AGG GTT AGG G |
| **HT-TT** | FAM-Human Telomere - TT spacer | /56-FAM/TTG GGT TAG GGT TAG GGT TAG GG |
| **HT-TTA** | FAM-Human Telomere - TTA spacer | /56-FAM/TTA GGG TTA GGG TTA GGG TTA GGG |
| **EHT** | FAM - Expanded Human Telomere | /56-FAM/TGG GGT TAG GGG TTA GGG GTT AGG GG |
| **TBA** | FAM-Thrombin Binding Aptamer | /56-FAM/GGT TGG TGT GGT TGG |
| **TBA-T** | FAM-Thrombin Binding Aptamer - T spacer | /56-FAM/TGG TTG GTG TGG TTG G |
| **TBA-C** | FAM-Thrombin Binding Aptamer - C spacer | /56-FAM/CGG TTG GTG TGG TTG G |
| **TBA-A** | FAM-Thrombin Binding Aptamer - A spacer | /56-FAM/AGG TTG GTG TGG TTG G |
| **TBA-TT** | FAM-Thrombin Binding Aptamer - TT spacer | /56-FAM/TTG GTT GGT GTG GTT GG |
| **TBA-TTA** | FAM-Thrombin Binding Aptamer - TTA spacer | /56-FAM/TTA GGT TGG TGT GGT TGG |
| **ETBA** | FAM - Expanded Thrombin Binding Aptamer | /56-FAM/TGG GTT GGG TGT GGG TTG GG |
| **Non-G4** | FAM-Duplex Top Strand | /56-FAM/CGA CTA TGA GGA TCT C |
| **HT-2** | Human Telomere 2 | /56-FAM/TG GGT TAG GGT TAG GGT TAG GGT T |
| **HT-2-I** | Human Telomere 2 Isomer | /56-FAM/TG TGT GAG TGT GAG TGT GAG TGT G |

### Supplementary Table 2. Fluorescence melting temperatures of oligonucleotides

| **Oligonucleotide** | **Buffer** | **H1** | **C1** | **H2** | **C2** | **Mean** | **Heat Mean** | **Cool Mean** |
| --- | --- | --- | --- | --- | --- | --- | --- | --- |
| **FAM-G4-IB** | Sodium | 53.7 | 52.3 | 53.9 | 52.6 | 53.1 | 53.8 | 52.5 |
| **FAM-G4-IB** | Potassium | 56.8 | 55.5 | 57.1 | 55.7 | 56.2 | 56.9 | 55.6 |
| **FAM-G4** | Sodium | 36.2 | 38.0 | 40.3 | 39.4 | 38.7 | 38.3 | 38.7 |
| **FAM-G4** | Potassium | 38.7 | 35.0 | 38.4 | 35.1 | 36.8 | 38.6 | 35.1 |
| **HT** | Sodium | 41.8 | 41.3 | 41.2 | 40.2 | 41.2 | 41.5 | 40.8 |
| **HT** | Potassium | 46.9 | 48.9 | 49.9 | 48.7 | 48.8 | 48.4 | 48.8 |
| **HT-T** | Sodium | 40.1 | 39.3 | 40.5 | 39.3 | 39.7 | 40.3 | 39.3 |
| **HT-T** | Potassium | 49.0 | 46.7 | 48.8 | 47.5 | 48.1 | 48.9 | 47.1 |
| **HT-C** | Sodium | 37.6 | 36.6 | 37.7 | 37.1 | 37.3 | 37.6 | 36.8 |
| **HT-C** | Potassium | 46.4 | 45.9 | 48.1 | 46.9 | 46.7 | 47.3 | 46.4 |
| **HT-A** | Sodium | 38.7 | 37.6 | 38.9 | 38.0 | 38.3 | 38.8 | 37.8 |
| **HT-A** | Potassium | 45.5 | 47.2 | 48.1 | 45.5 | 46.4 | 46.8 | 46.4 |
| **HT-TT** | Sodium | 39.9 | 39.0 | 40.3 | 39.1 | 39.5 | 40.1 | 39.1 |
| **HT-TT** | Potassium | 48.8 | 48.3 | 50.2 | 52.0 | 49.5 | 49.5 | 50.1 |
| **HT-TTA** | Sodium | 38.5 | 38.7 | 38.7 | 42.1 | 38.7 | 38.6 | 40.4 |
| **HT-TTA** | Potassium | 46.6 | 46.2 | 48.0 | 46.4 | 46.5 | 47.3 | 46.3 |
| **TBA** | Potassium | 28.3 | 27.4 | 28.2 | 27.7 | 28.0 | 28.2 | 27.5 |
| **TBA-T** | Potassium | 26.8 | 26.6 | 27.2 | 26.7 | 26.7 | 27.0 | 26.6 |
| **TBA-C** | Potassium | 27.1 | 25.7 | 27.3 | 26.0 | 26.5 | 27.2 | 25.8 |
| **TBA-A** | Potassium | 29.3 | 27.9 | 29.8 | 27.8 | 28.6 | 29.6 | 27.9 |
| **TBA-TT** | Potassium | 32.9 | 31.9 | 33.4 | 32.3 | 32.6 | 33.2 | 32.1 |
| **TBA-TTA** | Potassium | 47.3 | 42.5 | 52.2 | 42.5 | 44.9 | 49.7 | 42.5 |
| **EHT** | Sodium | 65.3 | 63.4 | 65.4 | 63.7 | 64.5 | 65.4 | 63.5 |
| **EHT** | Potassium | 82.2 | 74.8 | 84.5 | 75.4 | 78.8 | 83.3 | 75.1 |
| **ETBA** | Sodium | 38.3 | 37.1 | 38.8 | 37.5 | 37.9 | 38.5 | 37.3 |
| **ETBA** | Potassium | 55.7 | 56.2 | 56.6 | 54.0 | 56.0 | 56.1 | 55.1 |
| **HT-2** | Sodium | 33.9 | 32.8 | 34.2 | 33.0 | 33.5 | 34.0 | 32.9 |
| **HT-2** | Potassium | 45.9 | 43.9 | 46.5 | 43.9 | 44.9 | 46.2 | 43.9 |

### Supplementary Table 3. Absorbance melting temperatures of oligonucleotides

| **Oligonucleotide** | **Buffer** | **H1** | **C1** | **H2** | **C2** | **Mean** | **Heat Mean** | **Cool Mean** |
| --- | --- | --- | --- | --- | --- | --- | --- | --- |
| **FAM-G4-IB** | Sodium | 47.9 | 47.6 | 48.0 | 47.5 | 47.7 | 47.9 | 47.6 |
| **FAM-G4-IB** | Potassium | 51.4 | 50.9 | 51.7 | 50.7 | 51.1 | 51.6 | 50.8 |
| **FAM-G4** | Sodium | 36.8 | 36.4 | 36.7 | 36.4 | 36.6 | 36.8 | 36.4 |
| **FAM-G4** | Potassium | 39.8 | 38.4 | 39.1 | 38.3 | 38.8 | 39.5 | 38.4 |
| **HT** | Sodium | 41.2 | 40.5 | 40.9 | 40.6 | 40.8 | 41.1 | 40.6 |
| **HT** | Potassium | 48.9 | 48.1 | 48.4 | 48.2 | 48.3 | 48.6 | 48.2 |
| **HT-T** | Sodium | 42.0 | 41.4 | 41.7 | 41.6 | 41.6 | 41.8 | 41.5 |
| **HT-T** | Potassium | 52.1 | 51.8 | 52.0 | 52.0 | 52 | 52 | 51.9 |
| **HT-C** | Sodium | 39.2 | 38.8 | 39.4 | 38.8 | 39 | 39.3 | 38.8 |
| **HT-C** | Potassium | 48.6 | 47.9 | 48.7 | 47.8 | 48.2 | 48.7 | 47.8 |
| **HT-A** | Sodium | 39.2 | 39.1 | 39.3 | 39.1 | 39.1 | 39.3 | 39.1 |
| **HT-A** | Potassium | 49.0 | 48.4 | 49.1 | 48.3 | 48.7 | 49 | 48.4 |
| **HT-TT** | Sodium | 40.2 | 39.4 | 40.2 | 39.4 | 39.8 | 40.2 | 39.4 |
| **HT-TT** | Potassium | 50.6 | 49.9 | 50.7 | 49.7 | 50.2 | 50.6 | 49.8 |
| **HT-TTA** | Sodium | 38.5 | 38.7 | 38.7 | 42.1 | 38.7 | 38.6 | 40.4 |
| **HT-TTA** | Potassium | 45.8 | 45.3 | 45.9 | 45.3 | 45.5 | 45.9 | 45.3 |
| **TBA** | Potassium | 30.8 | 30.4 | 30.7 | 30.2 | 30.6 | 30.8 | 30.3 |
| **TBA-T** | Potassium | 30.4 | 29.8 | 30.4 | 29.7 | 30.1 | 30.4 | 29.7 |
| **TBA-C** | Potassium | 29.2 | 28.5 | 29.1 | 28.5 | 28.8 | 29.1 | 28.5 |
| **TBA-A** | Potassium | 27.7 | 27.2 | 27.6 | 27.3 | 27.4 | 27.6 | 27.2 |
| **TBA-TT** | Potassium | 31.7 | 31.2 | 31.7 | 31.2 | 31.5 | 31.7 | 31.2 |
| **TBA-TTA** | Potassium | 49.1 | 48.6 | 49.2 | 48.4 | 48.8 | 49.2 | 48.5 |
| **EHT** | Sodium | 63.5 | 62.8 | 63.6 | 62.8 | 63.2 | 63.5 | 62.8 |
| **EHT** | Potassium | 82.9 | 74.9 | 83.2 | 74.9 | 78.9 | 83 | 74.9 |
| **ETBA** | Sodium | 37.7 | 37.7 | 38.1 | 37.6 | 37.7 | 37.9 | 37.7 |
| **ETBA** | Potassium | 57.6 | 57.4 | 57.9 | 57.5 | 57.5 | 57.7 | 57.4 |
| **HT-2** | Sodium | 37.7 | 36.9 | 37.3 | 36.8 | 37.1 | 37.5 | 36.8 |
| **HT-2** | Potassium | 47.9 | 46.3 | 47.7 | 46.3 | 47 | 47.8 | 46.3 |

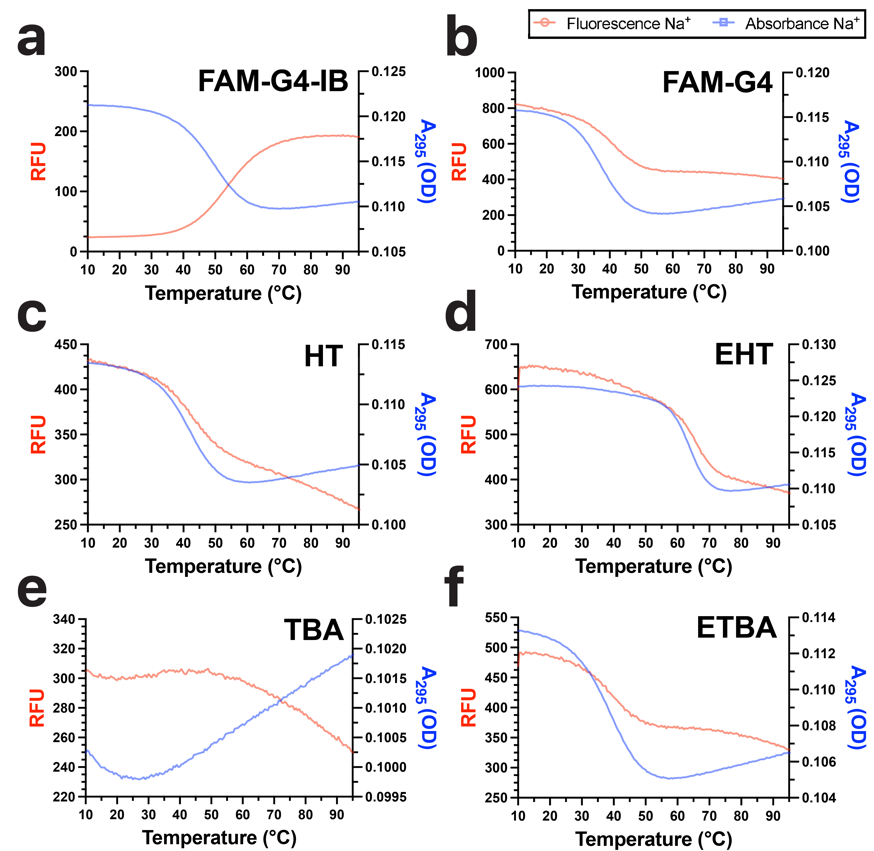

**Supplementary Figure 1. Quencher-free-fluorescence and absorbance-monitored thermal denaturation profiles both report on unfolding of various G-Quadruplex-forming sequences in sodium solution.** All samples in Na^+^ buffer. (a) **FAM-G4-IB** with fluorescein and Iowa Black quencher, (c) **FAM-G4** with only fluorescein modification, (d) **HT** with four repeats of human telomere sequence, (e) **EHT**, a variant of **HT** with an expanded G-tract. (f) **TBA**, the 15 nt thrombin binding aptamer, (g) **ETBA**, a variant of **TBA** with an expanded G-tract. Cooperative transitions were observed in all cases except that of **TBA**, indicating G-quadruplex to single-stranded transitions. The sodium form of **TBA** has sufficiently low stability that no transition was observed in sodium by both absorbance and quencher-free fluorescence.

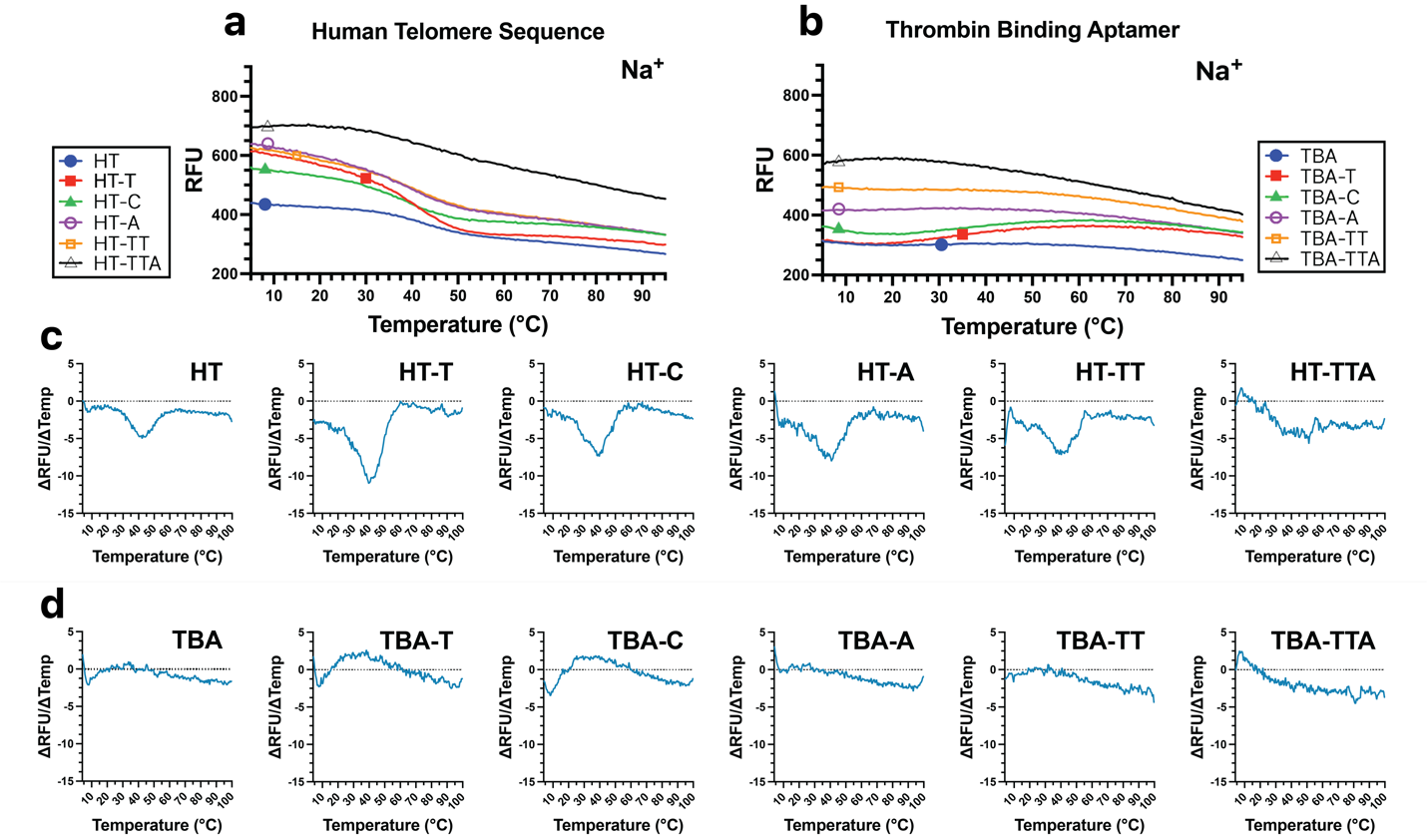

### Supplementary Figure 2. Influence of fluorophore proximity and spacer sequences on fluorescence transitions in two G-Quadruplexes in sodium solution.

(a) Fluorescence thermal melt of human telomere sequence (HT) with single or multiple-nucleotide spacers in Na^+^ buffer. (b) Fluorescence thermal melt of thrombin binding aptamer (TBA) with single or multiple-nucleotide spacers; no transition is observed due to low stability of this structure in sodium buffer. (c) First derivatives of HT fluorescence and temperature values in Na^+^ buffer (blue) with single or multiple nucleotide spacers (d) First derivatives of TBA fluorescence and temperature values in Na^+^ buffer (blue) with single or multiple nucleotide spacers; no transition is observed due to low stability of this structure in sodium buffer.

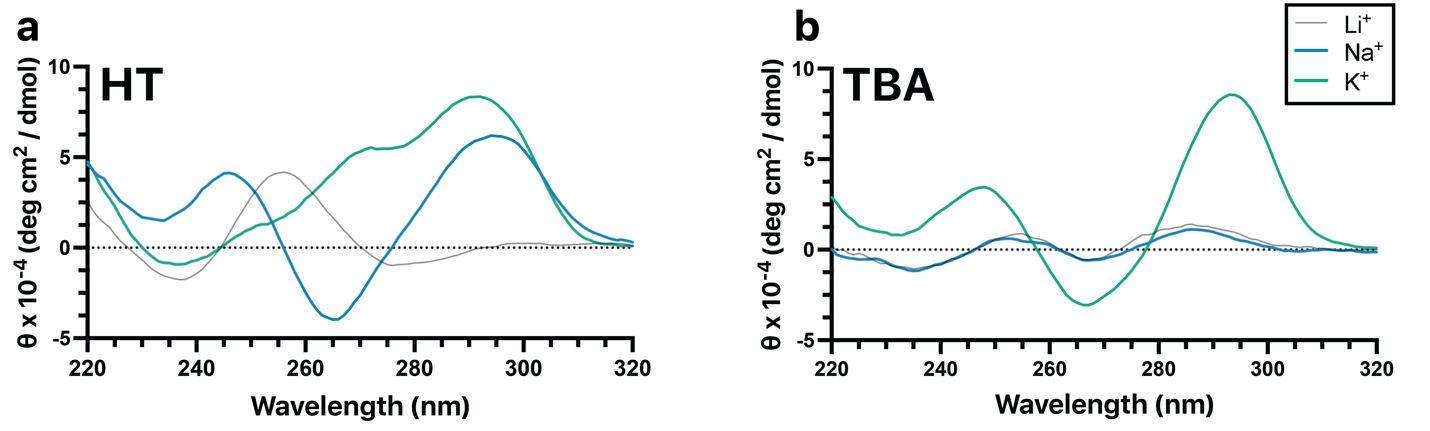

**Supplementary Figure 3. Circular dichroism spectra of G4-forming oligonucleotides.**

CD spectra measured between 220 nm to 320 nm at 25°C of a) **HT** are consistent with antiparallel (Na^+^) and hybrid (K^+^) topologies, while those for b) **TBA** are consistent with antiparallel topology in K^+^ solution and weak to no folding in Na^+^ solution.

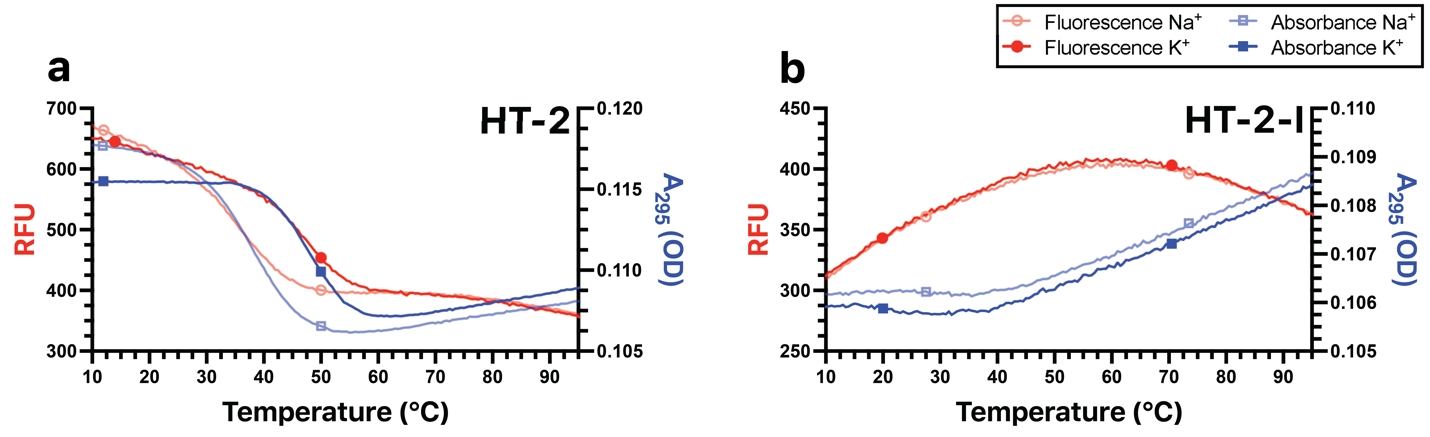

### Supplementary Figure 4. Experiments with two isomeric sequences illustrate that quencher-free fluorescence reports on secondary structure, rather than sequence content. (a) A second human telomere sequence HT-2 was employed gave cooperative unfolding transitions observed by quencher-free fluorescence and UV absorbance (A_295_). (b) An isomer of HT-2, HT-2-I, was examined by the same techniques. This isomer contained no consecutive guanosine residues, which was predicted to suppress G4 formation. As expected, HT-2-I gave no cooperative unfolding transition by quencher-free fluorescence or UV absorbance, illustrating that secondary structure formation, and not guanine content alone, was required to obtain the quencher-free fluorescence results observed.
